# PARPi triggers STING-dependent immune response and enhances therapeutic efficacy of immune checkpoint blockade independent of BRCAness

**DOI:** 10.1101/318980

**Authors:** Jianfeng Shen, Wei Zhao, Zhenlin Ju, Lulu Wang, Yang Peng, Marilyne Labrie, Timothy Yap, Gordon B. Mills, Guang Peng

## Abstract

Poly-(ADP-ribose) polymerase (PARP) inhibitors (PARPis) have shown remarkable therapeutic efficacy against *BRCA1/2* mutant cancers through a synthetic lethal interaction. PARPis are believed to exert their therapeutic effects mainly through the blockade of single-strand DNA damage repair, which leads to the accumulation of toxic DNA double strand breaks, specifically in cancer cells with DNA repair deficiency (BCRAness), including those harboring *BRCA1/2* mutations. Here, we show that PARPis modulate immune reposes, which contribute to their therapeutic effects independent of *BRCA1/2* mutations. The mechanism underlying this PARPi-induced reprogramming of anti-tumor microenvironment involves a promoted accumulation of cytosolic DNA fragments due to unresolved DNA lesions. This in turn activates the DNA sensing cGAS-STING pathway and stimulates production of type I interferons. Ultimately, these events promote PARPi-induced antitumor immunity independent of BRCAness, which can be further enhanced by immune checkpoint blockade. Our results may provide a mechanistic rationale for using PARPis as immunomodulatory agents to harness therapeutic efficacy of immune checkpoint blockade.

## Introduction

Poly (ADP-ribose) polymerase (PARP) inhibitors are approved for the treatment of patients with ovarian and breast cancers harboring *BRCA1* or *BRCA2* (*BRCA1/2*) mutations (1,2). The rationale supporting the development of single agent PARP inhibitors (PARPis) in *BRCA1/2* mutant cancers was based on the concept of synthetic lethality, which predicted antitumor efficacy in tumors with defects in homologous recombination (HR) repair, also known as BRCAness. PARP1 is the most abundant and ubiquitously expressed member of the PARP family and contributes the majority of PARP enzymatic activity and is thus the major target of PARPis (3). In the presence of DNA damage, PARP1 rapidly binds to DNA strand breaks and is essential for the repair of DNA single-strand breaks (SSBs) through base excision repair. In normal cells, recombinogenic DNA substrates generated by PARPis can be functionally resolved by the error-free HR repair pathway. In contrast, in cancer cells with defective HR repair, such as those deficient in BRCA1 or BRCA2, the DNA substrates generated by PARPis cannot be resolved, and therefore the cells are hypersensitive to PARPis (4,5).

Clinical studies have now also shown patient benefit with PARPis in those with *BRCA1/2* wildtype tumors (6). A recent Phase III clinical trial confirmed that patients with platinum-sensitive, recurrent ovarian cancer receiving PARPi treatment as maintenance therapy had significantly longer progression-free survival than those on placebo, regardless of *BRCA1/2* mutation status or HR repair status (7). These clinical observations raise the key question of whether PARPis can exert antitumor effects through mechanisms other than those leading to unresolved genomic lesions in tumors with DNA repair deficiency.

In this study, we show that PARPi treatment induces IFN-mediated antitumor immune responses. PARPis generate cytosolic double-strand DNA (dsDNA), which activate STING signaling and its associated-transcription programs. These critical changes amplify STING signaling and promote tumor infiltrating lymphocytes (TILs) and antitumor immunity, which can be further enhanced through immune checkpoint blockade.

## Materials and Methods

### Cell culture

Cell lines were validated by short tandem repeat (STR) DNA fingerprinting using the AmpF STR identifier kit according to the manufacturer’s instructions (Applied Biosystems, catalogue no. 4322288). The STR profiles were compared to known American Type Culture Collection fingerprints; to the Cell Line Integrated Molecular Authentication database, version 0.1.200808 (*Nucleic Acids Research* 2009; 37:D925-D932); and to the MD Anderson fingerprint database. The STR profiles matched known DNA fingerprints or were unique. The colorectal and ovarian cancer cell lines were kindly provided by Dr. Gordon B. Mills’ laboratory at The University of Texas MD Anderson Cancer Center. Cell line authentication was performed in the MD Anderson Characterized Cell Line Core in 2012 and 2013. All media were supplemented with 10% FBS with glutamine, penicillin, and streptomycin. The ID8 mouse ovarian surface epithelial cells were kindly provided by Dr. Vahid Afshar-Kharghan’s laboratory at MD Anderson. The ID8 cells were maintained in DMEM (high-glucose, Cellgro) supplemented with 4% FBS, 100 U/ml penicillin, 100 μg/ml streptomycin, 5 μg/ml insulin, 5 μg/ml transferrin, and 5 ng/ml sodium selenite. Cells were incubated at 37°C in a humidified incubator with 5% CO_2_.

### Antibodies and reagents

Anti-γH2AX (JBW301) antibodies were purchase from Millipore Sigma. Anti-β-Actin (A2228), anti-α-Tubulin (T6074) and anti-γ-Tubulin (SAB4503045) antibodies were purchase from Sigma-Aldrich. Anti-STING (D2P2F, #13647), anti-cGAS (D1D3G, #15102), anti-IRF3 (D6I4C, #11904), anti-phospho-IRF3 (Ser396, D6O1M, #29047), anti-TBK1 (D1B4, #3504), anti-phospho-TBK1 (Ser172, D52C2, #5483), anti-CtIP (D76F7, #9201), anti-MRE11 (#4895) and anti-PD-L1 (13684 and 64988) antibodies were purchased from Cell Signaling Technology. Anti-CD8 (sc-7970), anti-BLM (B-4, sc-365753), anti-EXO1 (SPM394, sc-56387) and other antibodies were purchased from Santa Cruz Biotechnology. BMN673 (S7048, Talazoparib) was purchased from Selleck Chemicals. VivoGlo Luciferin was purchased from Promega. Isotype control IgG and anti-PD-L1 (BE0101, clone 10F.9G2) antibodies were purchased from Bio X Cell. The ELISA kits of CCL5 and CXCL10 were purchase from Thermofisher. The multiplexed immunofluorescence IHC kit was purchased from PerkinElmer.

### RNA interference

Knockdown was achieved by RNA interference using a lentiviral vector–based MISSION shRNA (Sigma-Aldrich). The shRNA sequences were as follows: mouse *Sting* (NM_028261), TRCN0000346319 (#1), AGAGGTCACCGCTCCAAATAT; TRCN0000346266 (#2), CAACATTCGATTCCGAGATAT. SMART pool ON-Target plus siRNA for CtIP (L-011376-00), BLM (L-007287-00), EXO1 (L-007287-00), STING (L-024333-02), IRF3 (L-024333-02), TBK1 (L-024333-02) and cGAS (L-015607-02) were purchased from GE Dharmacon. Specificity and efficacy of knockdown was evaluated by Western blotting.

### Immunoblotting and Immunofluorescence

Cells were washed in PBS, and cellular proteins were extracted in 8 mol/L urea lysis buffer plus protease and phosphatase inhibitors (GenDEPOT) for 30 min at 4°C. Lysates were cleared by centrifugation, and proteins were separated by gel electrophoresis. Membranes were blocked in PBS 0.1% Tween 20 (PBS-T)/5% (w/v) milk for 1 hr at room temperature. Membranes were then incubated with primary antibodies diluted in PBS-T/5% (w/v) milk at 4°C overnight. Subsequently, membranes were washed with PBS-T and incubated with horseradish peroxidase secondary antibody (1:2,000; Jackson ImmunoResearch) diluted in PBS-T/5% skim milk. Membranes were washed in PBS-T, and bound antibody was detected by enhanced chemiluminescence (GE Healthcare). For detection of subcellular localization of IRF3, phospho-IRF3, TBK1 and phospho-TBK1, immunofluorescent staining was performed essentially as described previously (8). After treatment, cells were first fixed in ice-cold methanol for 10 min at −20°C, then blocked with 10% goat serum for 30 min at room temperature. Primary antibodies (IRF3, 1:200; Phospho-IRF3, 1:200; TBK1, 1:200; phospho-TBK1, 1:100, Cell Signaling Technology) were incubated at 4°C overnight, and Alexa 488– or Alexa 594–conjugated secondary antibodies (1:500,Thermo Fisher Scientific) were incubated for 1 hr at room temperature. Slides were mounted in ProLong anti-fade mounting medium containing DAPI (Thermo Fisher Scientific) and analyzed under a fluorescence microscope. At least 50 cells per sample were analyzed, and the percentage of cells with positive staining was determined.

### PicoGreen staining

PicoGreen staining was performed using Quant-iT Pico-Green dsDNA reagent and kits from Thermo Fisher Scientific. For confocal microscopy, PicoGreen was diluted into cell culture medium at the concentration of 3μl/ml, and the cells were incubated in the presence of PicoGreen at 37 °C for 1 h. The cells were washed and fixed for confocal microscopy with DAPI counterstaining.

### Multiplexed IHC staining

Tumor tissue retrieved from ID8 i.p. injection or CT26 subcutaneous injection were subjected to fixation and paraffin embedding. The sections cut from paraffin blocks were baked at 60 °C for 1 hour and deparaffinized and rehydrated with serial passage through changes of xylene and graded alcohol and washed in water. Multiplexed immunofluorescence was performed following the manufacturer’s instruction (PerkinElmer). The following antibodies were used for IHC: anti-mouse PD-L1 (D5V3B, Cell Signaling Technology, 1:100), anti-CD8 (H160, Santa Cruz, 1:200), anti-STING (D2P2F, Cell Signaling Technology, 1:100) and anti-phospho-IRF3 (D6O1M, Cell Signaling Technology, 1:100). Stained slides were counterstained with DAPI and coverslipped for review. Positivity was defined as≥5% of staining or the percentage of positive cells per slide was calculated.

### Quantitative PCR (Q-PCR)

Total RNA (1-2 μg) was used in a reverse transcriptase reaction with the High-Capacity RNA-to-cDNA Kit (Thermo Fisher Scientific). The SYBR Green Real-Time PCR Master Mixes kit (Life Technologies) was used for the thermocycling reaction in an ABI-VIIA7 RealTime PCR machine (Applied Biosystems). The Q-PCR analysis was carried out in triplicate with the following primer sets: mouse *Ccl5* (Forward: 5’-ATATGGCTCGGACACCACTC-3’; Reverse: 5’-TCCTTCGAGTGACAAACACG-3’), mouse *Cxcl10* (Forward: 5’- CCCACGTGTTGAGATCATTG-3’; Reverse: 5’-GTGTGTGCGTGGCTTCACT-3’), mouse *Gapdh* (Forward: 5’- ACCCAGAAGACTGTGGATGG-3’; Reverse: 5’- ACACATTGGGGGTAGGAAC-3’), human *CCL5* (Forward: 5’- TGCCCACATCAAGGAGTATTT-3’; Reverse: 5’- CTTTCGGGTGACAAAGACG-3’), human *CXCL10* (Forward: 5’- GGCCATCAAGAATTTACTGAAAGCA-3’; Reverse: 5’- TCTGTGTGGTCCATCCTTGGAA-3’), and human β-Actin (Forward: 5’- GAGCACAGAGCCTCGCCTTT-3’; Reverse: 5’ TCATCATCCATGGTGAGCTG-3’)

### ELISA

The cell culture supernatant or ascites from ID8 model were collected and processed according to the manufacturer’s instructions. The CXCL10 and CCL5 levels were determined using ELISA kits from R&D/Thermo Fisher Scientific following the standard procedures.

### *In vivo* mouse models

All studies were supervised and approved by the MD Anderson Institutional Animal Care and Use Committee (IACUC). Female mice were used as models to study ovarian cancer. When used in a power calculation, our sample size predetermination experiments indicated that 5 mice per group could identify the expected effects with 90% power. **Ovarian Cancer Syngeneic Model**: Luciferase labeled ID8 ovarian cancer cells (5 × 10^6^) were injected into th e peritoneal cavity of C57BL/6 mice per group (6–8 weeks old, CRL/NCI). The mice were allowed to recover and were monitored closely for the next 24 hours. Tumor progression was monitored once per week by Xenogen IVIS Spectrum *In Vivo* Bioluminescence Imaging System. Tumor volume was determined based on total flux (photons per second). Tumor-bearing mice were treated intraperitoneally (i.p.) with isotype control IgG or anti-PD-L1 antibody (200 μg/mouse, B7-H1, clone 10F.9G2, Bio X Cell) every three days. BMN673 was administered by daily oral gavage with a dose of 0.33 mg/kg. Mice reaching an endpoint requiring euthanasia by IACUC guidelines or weighing more than 35 grams as a result of tumor growth and/or ascites were euthanized. **Colorectal Cancer Syngeneic Model**: Murine CT26 colorectal cancer cells (2 × 10^5^) were subcutaneously injected into the left flank of BALB/C mice (6–8 weeks old, CRL/NCI) as previously described. Mice were allowed to recover and monitored closely for the next 24 hours. Tumor size was measured every three days and tumor volume was determined based on the calculation (width × width × length)/2. Tumor bearing mice were treated (i.p.) with isotype control IgG or anti-PD-L1 antibody (200 μg/mouse, B7-H1, clone 10F.9G2, Bio X Cell) every three days.

BMN673 was administered by daily oral gavage with a dose of 0.33 mg/kg. Mice reaching an endpoint requiring euthanasia by IACUC guidelines or exceeding tumor burden limits were euthanized. **Colorectal Cancer Nude Mouse Model**: Nude mouse experiments were conducted as described previously. Briefly, murine colorectal cancer cells CT26 (2 x 10^5^) were subcutaneously injected into the left flank of athymic nude mice (6–8 weeks old, CRL/NCI). Mice were allowed to recover and monitored closely for the next 24 hours. Tumor size was measured every three days and the tumor volume was determined based on the calculation (width × width × length)/2. Tumor bearing mice were treated (i.p.) with isotype control IgG or anti-PD-L1 antibody (200 μg/mouse, B7-H1, clone 10F.9G2, Bio X Cell) every three days.

BMN673 was administered by daily oral gavage with a dose of 0.33 mg/kg. Mice reaching an endpoint requiring euthanasia by IACUC guidelines or exceeding tumor burden limits were euthanized. **Ovarian Cancer Nude Mouse Model**: Luciferase labeled ID8 cells (5 x 10^6^) were injected into peritoneal cavity of athymic nude mice (6–8 weeks old, CRL/NCI). Mice were allowed to recover and monitored closely for the next 24 hours. Tumor progression was monitored once a week by Xenogen IVIS Spectrum in vivo bioluminescence imaging system. Tumor volume was determined based on total flux (photons per second). Tumor bearing mice were treated (i.p.) with isotype control IgG or anti-PD-L1 antibody (200 μg/mouse, B7-H1, clone 10F.9G2, Bio X Cell) every three days. BMN673 were administered by daily oral gavage with a dose of 0.33 mg/kg. Mice reaching an endpoint requiring euthanasia by IACUC guidelines or weighing more than 35 grams as a result of tumor growth and/or ascites were euthanized.

### Statistics

All statistical analyses were done in GraphPad Prism 7 software. Overall survival of various treatment groups was analyzed using the Cox regression model. Otherwise, unpaired t-tests were used to generate two-tailed P values.

## Results

### PARPi induces an accumulation of cytosolic DNA and activates STING signaling pathway

PARPi treatment markedly induced DNA double-strand breaks (DSBs) as detected by increased γ-H2AX levels, and thus caused cell cycle arrest in S phase (**Supplementary Fig. S1A and S1B**). The cytosolic DNA sensor cGAS is the most potent activator of the STING signaling pathway (9). After the recognition of cytosolic DNA, cGAS activates STING via generation of 2’-5’ cyclic GMP-AMP (cGAMP). STING, in turn, induces phosphorylation and nuclear translocation of IFN transcriptional regulatory factors TANK-binding kinase 1 (TBK1) and IFN regulatory factor 3 (IRF3) (10,11). We thus sought to determine whether PARPi induces accumulation of cytosolic DNA that could activate the cGAS-STING-TBK1-IRF3 axis in ovarian cancer cell lines HOC1 (*BRCA1/2* WT), UPN251 (*BRCA1* deleterious and restoration mutations, functional WT) (12) and HeLa (*BRCA1/2* WT).

As indicated by staining with the dsDNA-specific dye PicoGreen, BMN673 caused a significant accumulation of cytosolic dsDNA in multiple cell lines (**Fig. 1A)**. Moreover, phosphorylation of IRF3 and TBK1, two key components along the STING signaling pathway, was markedly elevated by BMN673 treatment in a time-dependent manner in three different cancer cell lines (**Fig. 1B**). Immunofluorescent staining also showed that PARPi remarkably induced the translocation of phospho-IRF3, phospho-TKB1, as well as total IRF3 and total TBK1 from the cytoplasm to the nucleus (**Fig. 1C**), which indicated functional activation of STING signaling. We then examined mRNA expression of *CCL5* and *CXCL10*, two major target genes downstream of STING activation that are involved in T-cell chemotaxis (13). We found a time-dependent increase in *CCL5* and *CXCL10* mRNA levels after PARPi treatment (**Fig. 1D**). Consistent with these changes in mRNA levels, PARPi substantially increased the production of CXCL10 as detected by ELISA (**Fig. 1E**). To determine if cGAS-STING-TBK1-IRF3 signaling is essential to the induction of chemokine expression by PARPi, we knocked down STING, TBK1, IRF3 or cGAS (**Supplementary Fig. S2A, S3A**) and treated cells with PARPi. We found significantly reduced upregulation of *CCL5* and *CXCL10* in response to PARPi treatment in these cells (**Fig. 1F and Supplementary Fig. S2B, S2C**). Together, these results demonstrate that PARPi induces accumulation of cytosolic dsDNA and activation of cGAS-STING-TBK1-IRF3 signaling to promote chemokine expression.

**Fig. 1.**
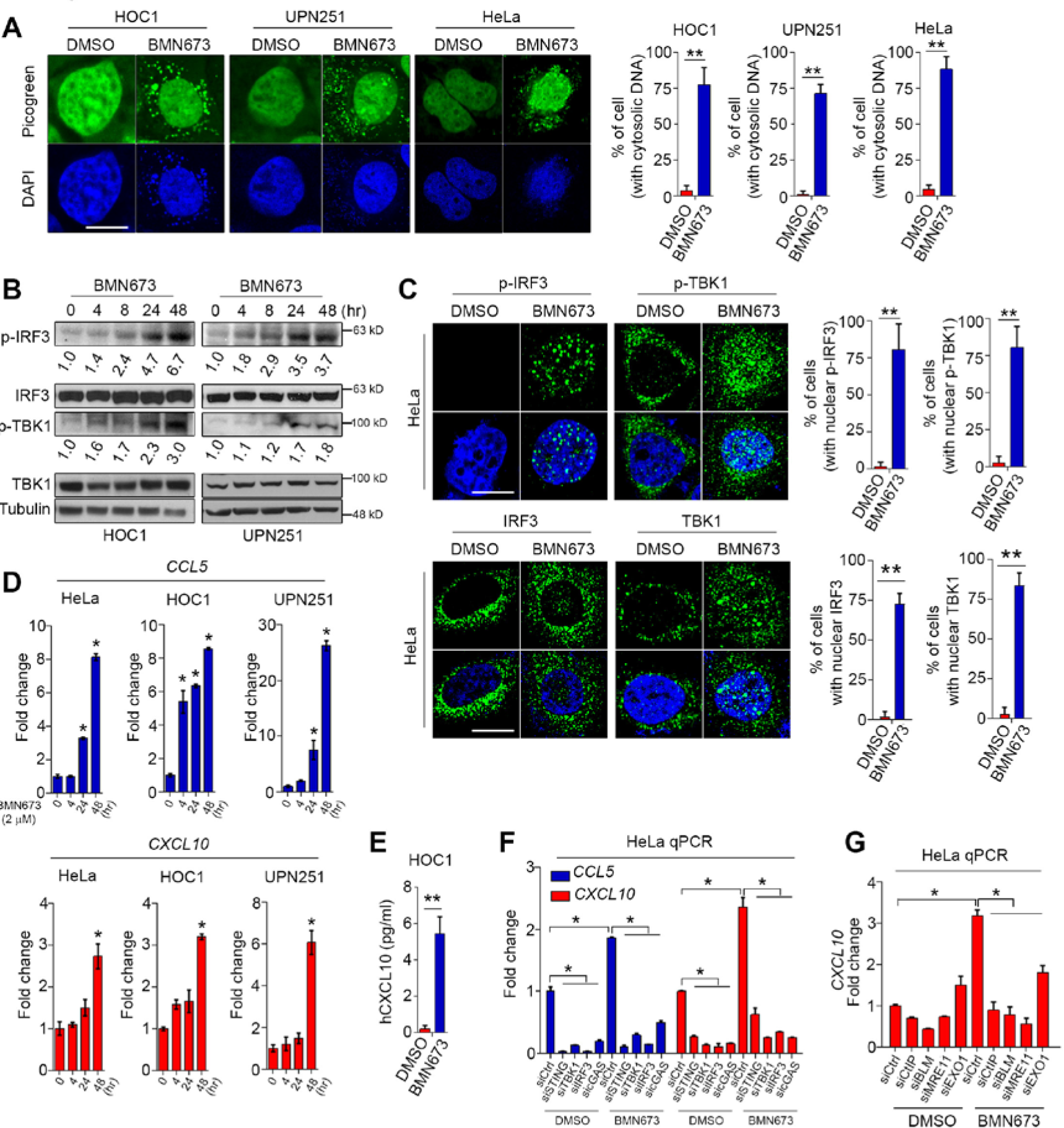
PARPi induces accumulation of cytosolic DNA and activates the STING signaling pathway. **A,** Representative images and quantitative analysis of PicoGreen staining in HOC1, UPN251, and HeLa cells treated with DMSO or BMN673 (2 μM) for 48 hours. DAPI (blue) was used to visualize the nuclei. Data represent mean ± s.e.m. of three independent experiments. **B**, Western blots of phosphorylated IRF3 (p-IRF3), total IRF3 (IRF3), phosphorylated TBK1 (p-TBK1) and total TBK1 (TBK1) in HOC1 and UPN251 cells treated with BMN673 (2 μM) for various times. The numbers represent the mean relative expression levels of p-IRF3 and p-TBK1 from three independent experiments. **C**, Representative immunofluorescent images and quantitative analysis of p-IRF3, p-TBK1, total IRF3 and total TBK1 staining in HeLa cell treated with DMSO or BMN673 (2 μM) for 48 hours. DAPI (blue) was used to visualize the nuclei. Data represent mean ± s.e.m. of three independent experiments. **D**, qPCR evaluation of *CCL5* and *CXCL10* expression in HeLa, HOC1, and UPN251 cells under DMSO or BMN673 treatment. Data represent mean ± s.e.m. of three independent experiments. **E**, ELISA quantification of human CXCL10 (hCXCL10) level in HOC1 cells under DMSO or BMN673 treatment. Data represent mean ± s.e.m. of three independent experiments. **F**, Quantitative PCR (qPCR) evaluation of *CCL5* and *CXCL10* levels in HeLa cells with depletion of STING (siSTING), TBK1 (siTBK1), IRF3 (siIRF3), or cGAS (sicGAS). Cells were treated with DMSO or BMN673 (2 μM) for 48 hours. Data represent mean ± s.e.m. of three independent experiments. **G**, qPCR evaluation of *CXCL10* level in HeLa cells with depletion of CtIP (siCtIP), BLM (siBLM), MRE11 (siMRE11), or EXO1 (siEXO1). Cells were treated with DMSO or BMN673 (2 μM) for 48 hours. Data represent mean ± s.e.m. of three independent experiments. Scale bar, 10 μM, ^*^, p<0.05; ^**^, p<0.01.

To identify the source of the cytosolic dsDNA induced by PARPi, we examined if factors involved in the degradation of DNA substrates at replication lesions might be required for the STING-dependent induction of immune response by PARPi. Several key factors containing and/or regulating nuclease activity, including MRE11, CtIP, BLM and EXO1, are recruited to DSBs, which can produce DNA fragments during HR repair and maintenance of replication fork stability (14). We reasoned that trapping of PARP1 by PARPi forms a barrier against DSB end resection and HR repair, which may lead to generation of dsDNA through degradation of unrepaired reversed replication forks. Indeed, we demonstrated that knockdown of these resection factors by siRNAs markedly reduced mRNA expression of *CXCL10* (**Fig. 1G and Supplementary Fig. S3B**), suggesting a key role for the requirement of these factors in generating PARPi-induced immune responses.

### PARPi activates STING signaling and immune checkpoint in vivo

We next investigated the *in vivo* effects of PARPi-induced immune responses. We used syngeneic immunocompetent mouse models of ovarian cancer (ID8) and colon cancer (CT26), which were treated with a clinically-relevant dosage of BMN673 (0.33 mg/kg). CT26 and ID8 cells have no known mutations in genes involved in the HR repair pathway, including *BRCA1/2* mutations. As expected, PARPi treatment exhibited no therapeutic effects in the xenograft model with the intraperitoneal injection of ID8 cells (**Fig. 2A** and **2B**). However, in the ID8 syngeneic model, PARPi can remarkably reduce tumor growth and significantly prolong survival (**Fig. 2C**). These data indicated that an intact T cell-mediated immune response is required for PARPi efficacy in the ID8 model.

**Fig. 2.**
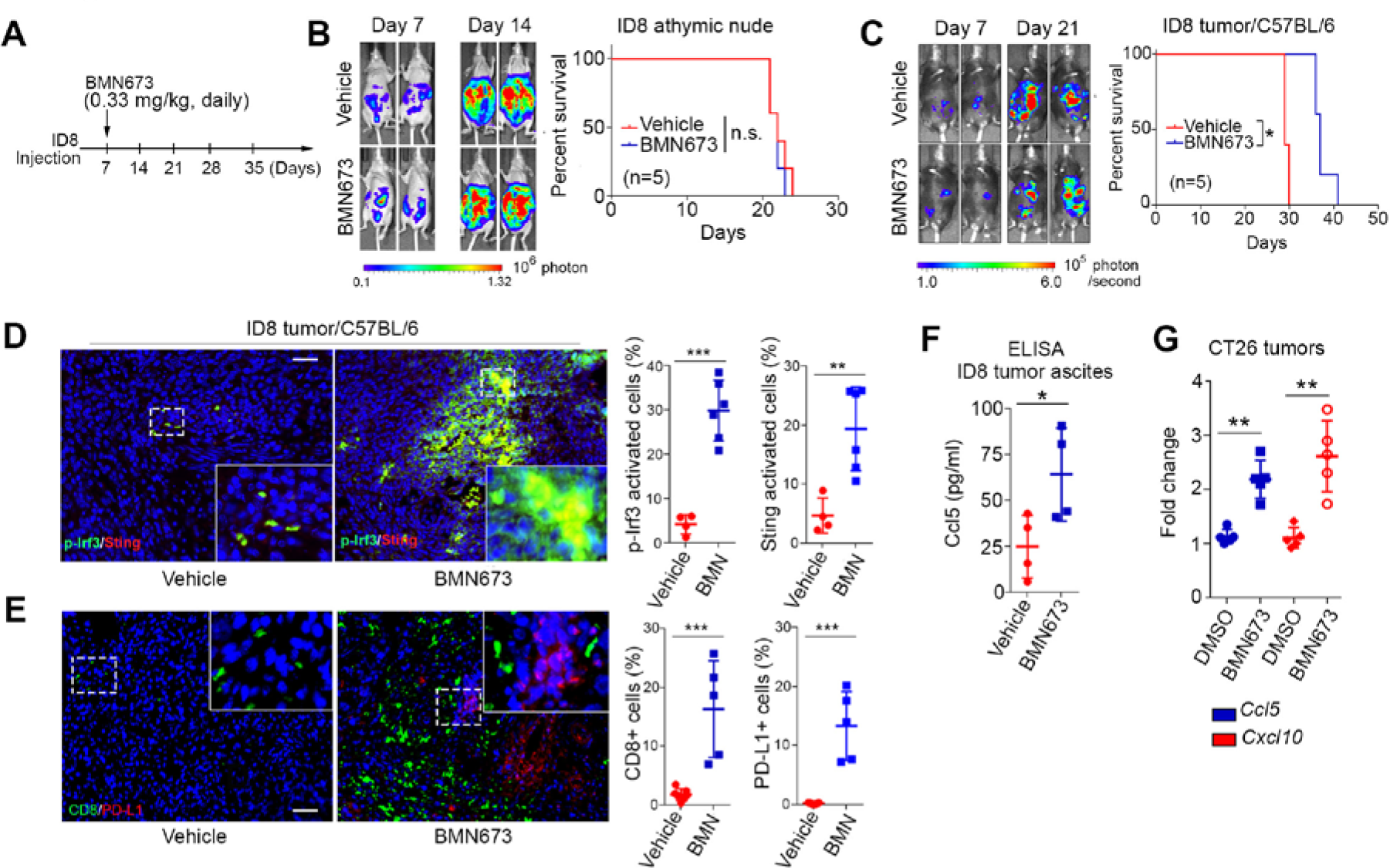
PARPi activates STING signaling and immune checkpoint *in vivo*. **A,** Schematic of PARPi treatment in intraperitoneal xenograft and syngeneic ID8 mouse models. BMN673 (0.33 mg/kg) was orally administered daily. Treatment was started on day 7 after ID8 cell inoculation and continued until the mice were euthanized (n=5). **B** and **C**, Representative bioluminescence images (left panels) of mice with intraperitoneal (i.p.) xenograft tumors (**B**) and syngeneic tumors (**C**) 7 and 21 days after ID8 cell inoculation and survival curves (right panels) of mice with ID8 i.p. xenograft tumors (**B**) and syngeneic tumors (**C**). ^*^, p<0.05; n.s. not significant. **D**, Representative images and quantitative analysis of phosphorylated Irf3 (p-Irf3) and Sting in ID8 tumors from C57BL/6 mice 30 days after tumor cell inoculation. DAPI (blue) was used to visualize the nuclei. **E**, Representative images and quantitative analysis of Cd8a and Pd-l1 in ID8 tumors from C57BL/6 mice 30 days after tumor cell inoculation. DAPI (blue) was used to visualize the nuclei. **F**, ELISA evaluation of Ccl5 levels in ascites from C57BL/6 mice when euthanization was performed. **G**, qPCR of *Ccl5* and *Cxcl10*, and levels in CT26 tumors from BALB/C mice 22 days after tumor cell inoculation. Dashed square, area for magnification. Scale bar, 50 μM, ^*^, p<0.05; ^**^, p<0.01; ^***^, p<0.001; n.s. not significant.

It has been reported that expression levels of *CCL5* and *CXCL10* positively correlate with infiltrating CD8^+^ cytotoxic lymphocytes in various cancers (15,16). We thus conducted immunohistochemistry (IHC) analysis of STING activation and immune response. In syngeneic ID8 and CT26 models, PARPi significantly upregulated the levels of Sting and phospho-Irf3, indicating robust activation of the Sting signaling pathway *in vivo* (**Fig. 2D and Supplementary Fig. S4A**). Consistent with this finding, remarkably higher percentages of CD8 positivity, a key cytotoxic T lymphocyte marker (17), as well as higher percentages of PD-L1 positivity, a marker of immune checkpoint activation, were found in PARPi-treated tumors than in control tumors in both models (**Fig. 2E and Supplementary Fig. S4B**).

Furthermore, we used ELISA and qPCR assays to validate functional activation of the Sting signaling pathway in animal models. As shown in **Fig. 2F** and **2G**, PARPi treatment induced expression of Ccl5 and Cxcl10, which was consistent with *in vitro* studies (**Fig. 1**). Together, these data showed that PARPi treatment induces an immunogenic response through the activation of the STING pathway and enhancement of type I IFN response and tumor-infiltrating lymphocytes *in vivo*.

However, activation of the PD-1/PD-L1 immune checkpoint pathway may counterbalance the impact of active tumor-infiltrating lymphocytes and block elimination of tumor cells despite the immunogenic microenvironment-induced by PARPi. These results raised the possibility that the combination of immune checkpoint blockade and PARPi would synergistically limit tumor growth and prolong survival by strongly priming PARPi-induced immunogenic responses.

### Immune checkpoint blockade targeting PD-1/PD-L1 pathway potentiates therapeutic efficacy of PARPi in syngeneic mouse models

To test this possibility, we first treated mice with intraperitoneal ID8 tumors with control IgG, BMN673, anti-PD-L1, or the combination of BMN673 and anti-PD-L1 for 3 weeks and then stopped treatment and monitored tumor growth and mouse survival (**Fig. 3A**). We found that only the combination treatment significantly reduced tumor growth compared to IgG control while no significant changes in mouse weight were observed (**Fig. 3B-3E**). Consistent with relatively poor immunogenicity and low tumor-infiltrating lymphocyte levels in ID8 tumors (18), and the fact that the major role of the PD-1/PD-L1 immune checkpoint is to limit the activity of effector T cells in the tumor microenvironment, ID8 tumors did not significantly respond to anti-PD-L1 therapy after 3 weeks of treatment (**Fig. 3C-3E**). BMN673 alone reduced tumor growth; however, the reduction did not achieve statistical significance (**Fig. 3C-3E**). In the third week of treatment, mice developed ascites which interfered with further luciferase measurements, but the study was continued to measure survival. As shown in **Fig. 3F**, only the combination treatment significantly prolonged survival compared to the IgG control; treatment with single-agent BMN673 or anti-PD-L1 did not substantially alter survival. Continuous treatment with the combination of BMN673 and anti-PD-L1 produced a remarkably better outcome than treatment with this regimen for three weeks (**Fig. 3G**), consistent with ongoing activity of the combination regimen. Notably, continuous treatment with the combination therapy led to long-term survival in 2 of the 10 mice (20%) bearing ID8 tumors (**Fig. 3G**), suggesting a potential curative effect from this continuous regimen.

**Fig. 3.**
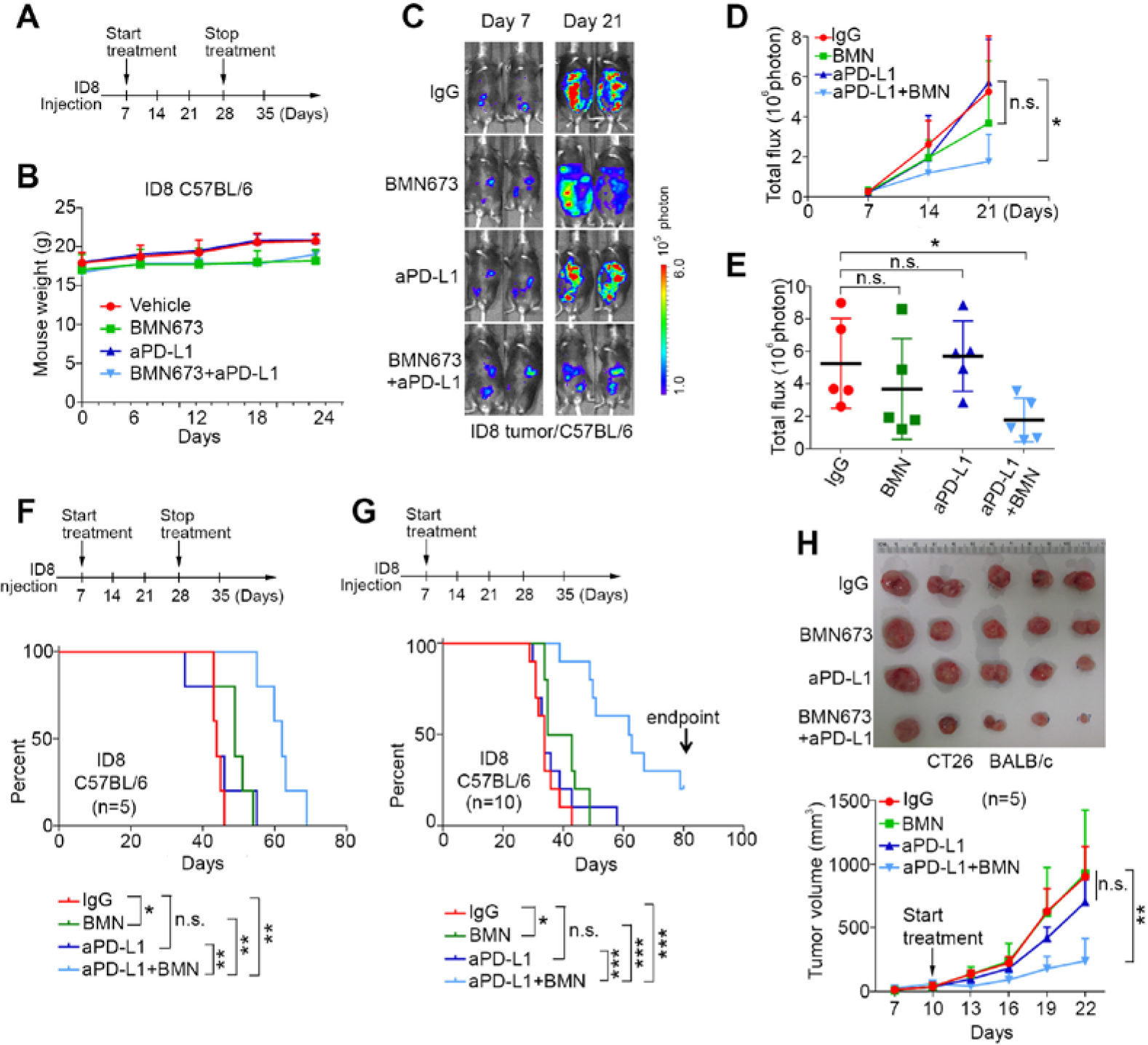
Immune checkpoint blockade targeting PD-1/PD-L1 pathway potentiates therapeutic efficacy of PARPi in syngenic mouse models. **A**, Schematic of treatment. Intraperitoneal (i.p.) injections of isotype control IgG (200 μg/mouse) and anti-PD-L1 antibody (aPD-L1, 200 μg/mouse) were started at day 7 and stopped at day 28 after ID8 cell inoculation. BMN673 (0.33 mg/kg) was orally administered daily. **B**, Weight of C57BL/6 mice bearing ID8 i.p. tumors and treated with vehicle, BMN673, anti-PD-L1 (aPD-L1), or BMN673 plus aPD-L1 over time (n=5). **C**, Representative bioluminescence images of mice bearing ID8 i.p. tumors after 7 and 21 days of inoculation. **D** and **E**, Statistical analysis of change in bioluminescence over time (mean ± s.e.m) (**D**) and bioluminescence at the end of the study (each dot represents one mouse) (**E**) (n=5). **F**, Survival curves of mice with ID8 i.p. tumors with treatment started on day 7 after ID8 cell inoculation and stopped on day 28, as indicated by the arrows (n=5). **G**, Survival curves of mice with ID8 i.p. tumors with treatment started on day 7 after ID8 cell inoculation and continued until the mice were euthanized (n=10). **H**, Representative images and tumor volume measurements of CT26 tumors in BALB/c mice with continuous treatment (n=5). ^*^, p<0.05; ^**^, p<0.01; ^***^, p<0.001; n.s. not significant.

We next used CT26 mouse colon tumor cells as an independent syngeneic model to validate the effects of combination therapy using BMN673 and anti-PD-L1. Because CT26 tumors grew aggressively, mice were treated for 12 days before the tumor size reached the maximum volume allowed by the animal protocol. Consistent with our observations in the ovarian cancer model, the combination therapy significantly reduced tumor burden compared to IgG control (**Fig. 3H**). However, these tumors were not sensitive to BMN673 alone. Collectively, these results indicated that the combination of BMN673 and anti-PD-L1 significantly inhibits tumor growth and prolongs survival in syngeneic cancer models.

### The therapeutic effects of combining PARPi and anti-PD-L1 depend on an intact immune system

To further confirm that the synergistic effects of the combination of BMN673 and anti-PD-L1 are dependent on an immune response, mouse xenografts of ID8 and CT26 were generated in nude mice, which do not contain a competent adaptive immune system. None of the treatments examined - BMN673 monotherapy, anti-PD-L1 monotherapy, and the combination of BMN673 and anti-PD-L1 - inhibited tumor growth or improved survival (**Fig. 4A-4D**). These results strongly supported the notion that an intact immune system is a prerequisite to achieve the benefits of combination therapy.

**Fig. 4.**
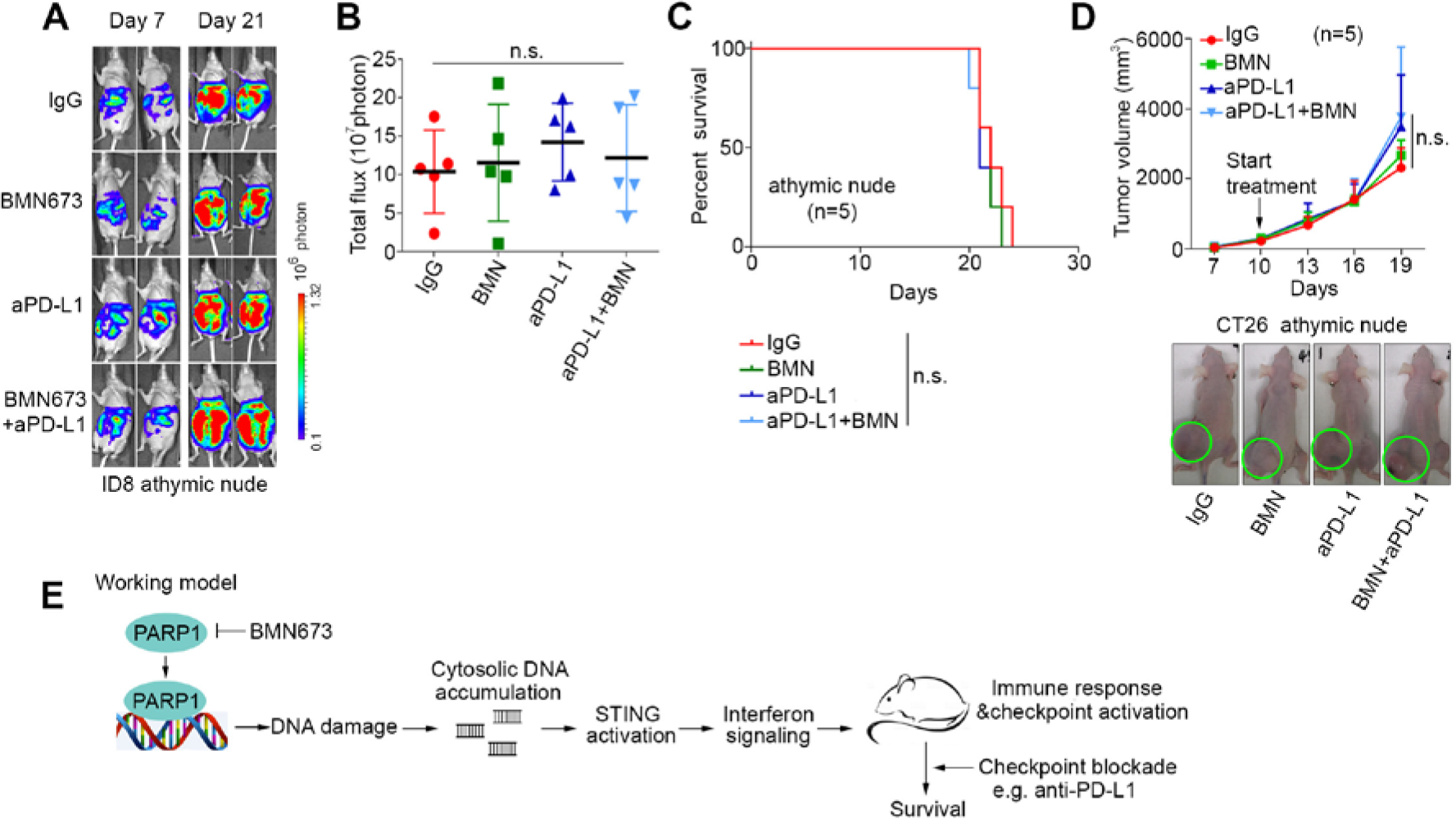
An intact immune system is required for the therapeutic benefits of combining PARPi with anti-PD-L1. **A,** Representative bioluminescence images of athymic nude mice bearing ID8 i.p. tumors. i.p. injection of isotype control IgG (200 μg/mouse) and anti-PD-L1 (aPD-L1, 200 μg/mouse) were stated at day 7 and continued until mice were euthanized. BMN673 (0.33 mg/kg) was orally administered daily. **B,** Statistical analysis of bioluminescence at the end of the study (n=5). Data represent mean ± s.e.m. **C**, Survival curves of athymic nude mice with ID8 i.p. tumors. Treatment was started on day 7 and continued until mice were euthanized. **D**, Representative images and tumor volume measurements of CT26 tumors in athymic nude mice with the indicated treatments (n=5). n.s., not significant. **E**, Model for the role of PARPis in inducing a distinct transcriptomic change through two interconnected STING-dependent mechanisms.

## Discussion

Here we show PARPi leads to an accumulation of cytosolic dsDNA and thereby activates the cGAS-STING-TBK1-IRF3 innate immune pathway, which upregulates transcriptional programs to produce type I IFN and its related immune responses **(Fig. 4E)**. Using two syngeneic mouse models with inherently less immunogenic tumors, we demonstrate that PARPi treatment enhances tumor susceptibility to immune checkpoint blockade. Importantly, these responses were observed regardless of the *BRCA1/2* mutation status of the cell lines assessed both in *vitro* and *in vivo*.

Recent studies have shown that micronuclei resulting from mis-segregation of DNA during cell division can be recognized by the cytosolic DNA sensor cyclic GMP-AMP synthase (cGAS) and activate the stimulator of interferon (IFN) genes (STING) innate immune pathway (19). However, it remains unknown whether PARPi-induced DNA damage is sufficient to generate cytosolic DNA that may serve as a signaling molecule to prime an immune response through the activation of the STING pathway. Our data showed that DNA damage induced by PARPi generates cytosolic DNA, primarily dsDNA, which functions as an immune stimulus engaging activation of the STING pathway. A recent study showed that ionizing radiation (IR) induces primarily cytosolic ssDNA (20). The different forms of cytosolic DNA resulting from DNA damage may depend on the mechanisms by which cells cope with DNA damage. In the presence of IR-induced DSB, DSB end resection is initiated for HR repair. ssDNA can be generated by nucleases that control DNA end resection including BLM and EXO1 (14). However, PARPis induces stalled or collapsed replication forks. dsDNA generated by PARPi may be related to DNA replication fork degradation/reversion or the restart of replication. Our study thus proposes a novel molecular mechanism underlying PARPi therapeutic effects, which is independent of its conventional cytotoxic effects resulting from unresolved DNA damage in DNA repair-deficient cancer cells.

Since the immunogenic responses induced by PARPi is not dependent on DNA repair deficiency, our findings may explain the antitumor activity observed in PARPi treated patients regardless of *BRCA1/2* mutation status or the presence of HR defective genomic signature (7). Our study also suggests that PARPi in combination with immune checkpoint inhibitors may demonstrate benefit in tumors regardless of DNA repair status, which has important clinical implications. Indeed, multiple Phase Ib combination trials, including BMN673 and the PD-L1 inhibitor avelumab are currently ongoing in the clinic. The results from the studies herein provide important information on mechanisms of sensitivity, as well as a route to potential biomarkers of efficacy.

